# The broad shell colour variation in common cockle (*Cerastoderma edule*) from Northeast Atlantic relies on a major QTL revealed by GWAS using a new high-density genetic map

**DOI:** 10.1101/2022.04.13.488192

**Authors:** Miguel Hermida, Diego Robledo, Seila Díaz, Damián Costas, Alicia L. Bruzos, Andrés Blanco, The Cockle’s Consortium, Paulino Martínez

## Abstract

Shell colour pattern shows broad diversity in molluscs, and both genetic and environmental factors seem to interact to some extent on the final phenotype. Despite information on the genetic component and pathways involved in shell construction and colour has increased in the last decade, more data are needed particularly to understand colour variation and its putative role on adaptation. The European common cockle (*Cerastoderma edule*) is a valuable species from ecological and commercial perspectives with important variation in colour pattern, but this diversity has never been characterized and the underlying genetic architecture is unknown. In this study, we constructed a high-density genetic map, as an essential tool for genomic screening in common cockle, that was applied to ascertain the genetic basis of colour pattern variation in the species. The consensus map, including 13,874 2b-RAD SNPs, was constituted by the 19 linkage groups (LGs) corresponding to the n = 19 chromosomes of its karyotype and spanned 1,073 cM (730 markers per LG; inter-marker distance of 0.13 cM). Five full-sib families showing segregation for several colour-associated traits were used to perform a GWAS analysis. A major QTL on chromosome 13 explained most of the variation for shell colour patterns. Mining on this genomic region revealed the presence of several candidate genes enriched on Gene Ontology terms such as anatomical structure development, ion transport, membrane transport and cell periphery, closely related to shell architecture, including six chitin-related, one ependymin, several ion binding and transporters, and others related to transit across the cell membrane. Interestingly, this major QTL overlaps with a genomic region previously reported associated with divergent selection in the distribution range of the species, suggesting a putative role on local adaptation.

## Introduction

The common cockle, *Cerastoderma edule*, is a bivalve mollusc naturally distributed along the north-eastern Atlantic coast, from Senegal in the South to Norway and Iceland in the North, inhabiting on intertidal soft sediment regions (Hayward & Ryland, 1995). This species has an important ecological role on marine sediment renewal and represents a food source for birds, crustaceans, and fish, thus playing an important role for coastal ecosystems and marine communities (Norris et al., 1998). The species is considered a delicacy and it is commercially fished mainly in Ireland, United Kingdom, Netherlands, France, Spain, and Portugal, where its commercialization represents a primary source for employment in coastal communities (http://www.cockles-project.eu/; Kamermans & Smaal, 2002).

As bivalve molluscs, the shell is a fundamental part of the cockle, serving as protection from predators, desiccation at intertidal zones or mechanical damage, enabling behaviours such as being swept by currents or burrowing. This multi-layered exoskeleton is formed mainly by calcium carbonate deposited into an organic matrix of proteins and pigments secreted by specialized epithelial cells on the dorsal mantle (Clark et al., 2020). Although a conserved set of regulatory genes appears to underlie mantle progenitor cell specification, the genes that contribute to the formation of the mature shell are diverse (McDougall and Degnan, 2018). Technical innovations have allowed to discover surprising patterns of shell pigmentation and rapid divergences in the mix of pigments used to achieve similar colour patterns (Miyamoto et al., 2013; Williams et al., 2017). Indeed, the shells of different bivalves are remarkably diverse, and shell pigmentation varies dramatically even within species. Variation in shell colour and pattern may be associated with different biotic or abiotic factors such as predation, substrate, diet, or environmental conditions (Williams, 2017; Vendrami et al., 2019). Since colour variation has been reported to be controlled to some extent by genetic factors, shell colour might be important for adaptation of bivalve population to selective pressures (Vendrami et al., 2017; Ding et al., 2021). Moreover, as a commercialized food resource, shell colour can be important for consumer pleasantness and acceptability, affecting to a certain extent the sale value (http://www.cockles-project.eu/). Whether common cockle populations show variation in their colouration patterns, if it has an underlying genetic basis and to what extent it could be related to adaptive variation is unknown.

The rapid expansion of next-generation sequencing (NGS) in the last decade has allowed the development of genotyping-by-sequencing (GBS) methods, which have been employed to discover and genotype thousands of single nucleotide polymorphisms (SNPs) in a cost-effective manner, enabling population-scale genetic studies in non-model species (Davey et al., 2011). These methods, including Restriction-site Associated DNA (RAD) sequencing (Baird et al., 2008) and its derivations, ddRAD (Peterson et al., 2012), 2b-RAD (Wang et al., 2012) or SLAF (Sun et al., 2013), have been successfully used for high-density genotyping in many aquaculture species (Robledo et al., 2018), including several important commercial molluscs (Gomes dos Santos et al., 2020). GBS methods have been recently applied to understand adaptive variation of common cockle from Northeast Atlantic, and consistent signals of adaptive variation were detected both at microgeographic (dd-RAD; Coscia et al., 2020) and macrogeographic (2b-RAD; Vera et al., 2022) scales. GBS have facilitated the construction of high-resolution linkage maps (Maroso et al., 2018; Dong et al., 2019; de la Herrán et al., 2022), which are important tools for genome scaffolding and assembly (Fierst, 2015), and have aided to disentangle the genetic basis of relevant evolutionary or productive traits through quantitative trait locus (QTL) screening (Aslam et al., 2020; Yin & Hedgecock, 2021). Genetic maps have been used to study the genetic architecture of traits of interest in various bivalve species, such as growth in Zhikong scallop (*Chlamys farreri*; Zhan et al., 2009), bay scallop (*Argopecten irradians*; Li et al., 2012) or *Crassostrea gigas* (Li et al., 2018), various pearl-quality traits in triangle sail mussel (*Hyriopsis cumingii*; Bai et al., 2016) and resistance to pathologies (Harrang et al., 2015). Previous studies have also identified QTL for shell colouration in several bivalves including Manila clam (*Ruditapes philippinarum*; Nie et al., 2017, 2020, 2021), hard clam (*Mercenaria mercenaria*; Hu et al., 2019), Pacific oyster (*Crassostrea gigas*; Feng et al., 2018; Song et al., 2018; Wang et al., 2018; Han et al., 2021), black-lip pearl oyster (*Pinctada margaritifera*; Lemer et al., 2015), Akoya pearl oyster (*Pinctada fucata martensii*; Xu et al., 2019) and Yesso scallop (*Patinopecten yessoensis;* Ding et al., 2015; Zhao et al., 2017), and therefore, similar strategies might be employed to ascertain the genetic component underlying differences in shell colouration in common cockle.

In this study, we investigated the variation of shell colouration patterns in Northeast Atlantic populations of European common cockle and studied the genetic architecture of this trait through GWAS on several full-sibs families using the first common cockle high-density linkage map here constructed using 2b-RAD SNP genotyping. A major QTL underlying colouration patterns in this species was detected on chromosome 13 and several candidate genes and enriched functions identified. This information should be considered as a potential source for adaptive variation in common cockle, and could be exploited in breeding programmes to adapt production to consumer demands.

## Material and methods

### Colour pattern variation in common cockle European populations

To assess the phenotypic diversity of cockle colour patterns across the Northeast Atlantic, several populations along its European distribution were studied. A total of 270 shells from nine cockle beds belonging to seven countries were provided by the Scuba Cancers Project (ERC-2016-STG) (Fig. 1).

**Figure 1.**
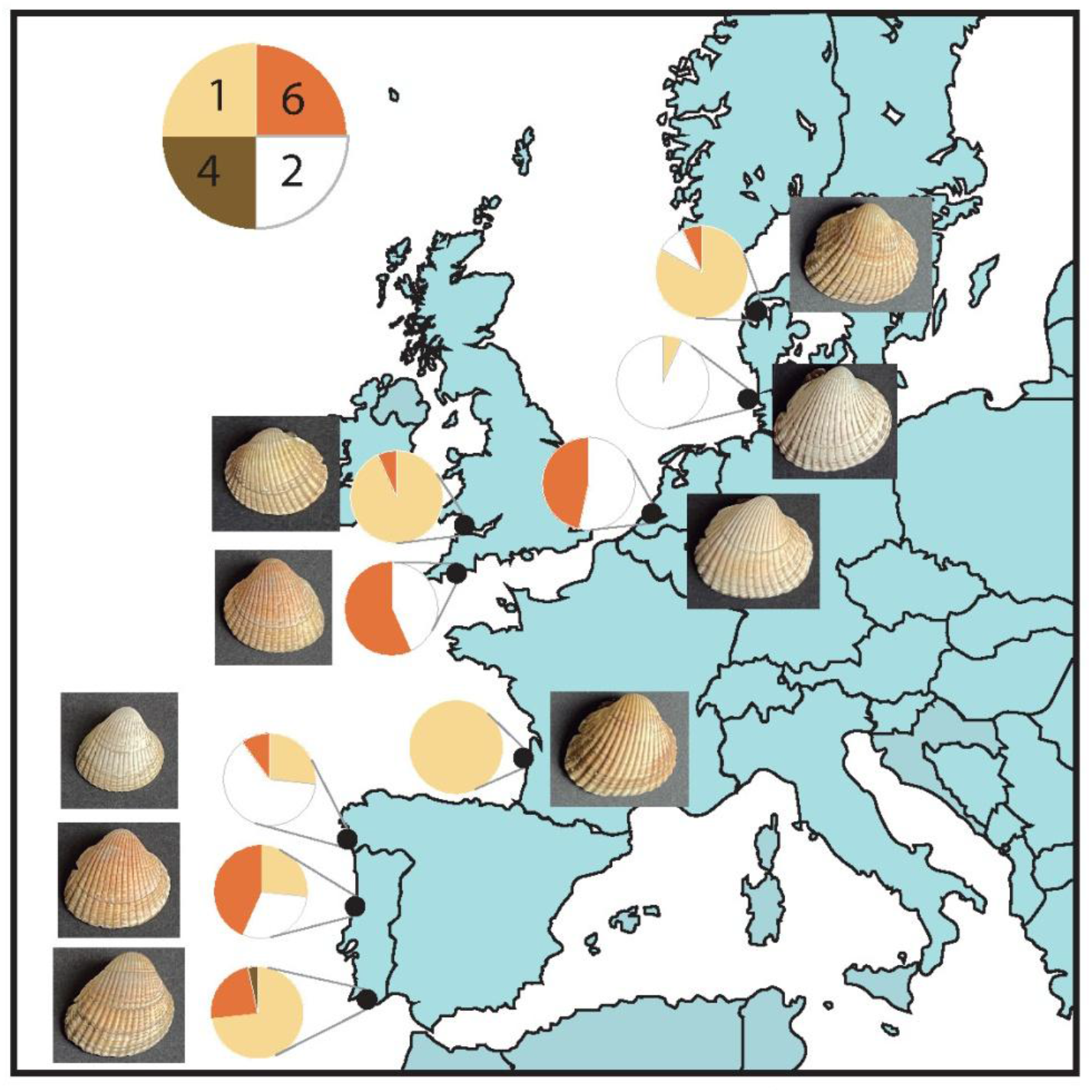
Geographical representation of the European samples of *C. edule*, including pie chart colour distributions (yellow (1), white (2), brown (4), orange (6)) and a photograph of a representative individual. From north to south: Nykobing Mors (Denmark), Sylt (Germany), Slikken van Viane (Netherlands), Wales and Plymouth (United Kingdom), Arcachon (France), Baiona (Spain), Aveiro and Algarve (Portugal).

The colour of each individual shell was classified as a categorical value, being numerically coded according to the most prevalent colour of the shell: yellow (1), white (2), grey (3), brown (4), black (5), and orange (6) (Fig. 2). Moreover, the presence of specific colour patterns of the shell was also recorded as three other traits: i) a circle in the umbo, differentiated from the rest of the shell and generally yellow (circle); ii) a broken white line (line); and iii) a stripe with white line edge (stripe) (Fig. 2).

**Figure 2.**
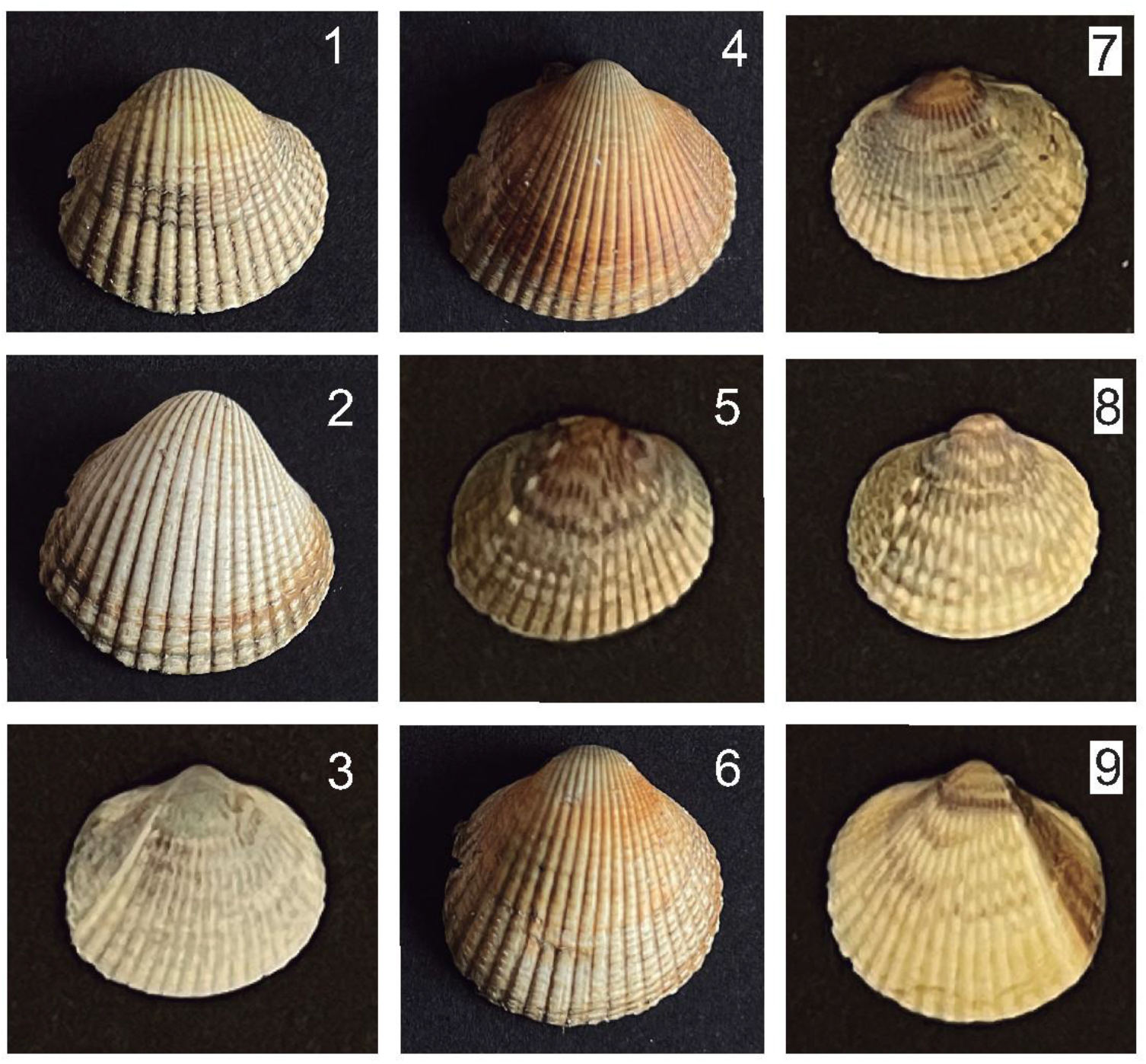
Representative individuals for each colour and pattern identified in shells of *C. edule* from Northeast Atlantic in this study. From up to down and left to right: colour phenotypes yellow (1), white (2), gray (3), brown (4), black (5), orange (6), and shell patterns circle (7), line (8) and stripe (9).

### Cockle families

In May 2018, 300 mature adult *C. edule* cockles were collected in Noia, Galicia (NW Spain) and transferred to the ECIMAT-CIM-UVigo marine facilities (Vigo, NW Spain). Cockles were kept individually in glasses with 0.3L of 1 μm filtered seawater at 20°C. Spawning was induced by thermal shocks between 10 and 22ºC for 10 hours and the quality of the oocytes and sperm was evaluated under a light microscope. Controlled fertilization was carried out by adding sperm to oocytes, one male x one female, at a ratio of 1:10. D-shaped larvae were obtained 24 hours after fertilization with a transformation rate from trochophore larvae of 42% ± 19. Following this protocol, a total of eleven full-sib families were obtained by crossing one male x one female, involving a total of 11 females and 7 males.

Larvae from each family were cultured in individual 150 L cylindrical-conical tanks at a density of 8 ± 3 larvae mL^−1^ with sea water filtered at 1 μm and treated with UV, slight aeration, temperature 19.0 ± 1.4 ºC, in an open circuit with a renewal of 5% volume / hour. The diet consisted of *Tisochrysis lutea* (ECC038), *Chaetoceros neogracile* (ECC007), *Phaeodactylum tricornutum* (ECC028) and *Rhodomonas lens* (ECC030) in a ratio of 1:1:1:1 (according to the cell count), and *Tetraselmis suecica* (ECC036) was included from the seventh day of culture. The daily diet was administered automatically every 4 hours in 6 daily intakes, maintaining a constant concentration in the tank of 20-40 cells μl^−1^. At 14 days post-fertilization (dpf), pediveliger larvae from each family were transferred to separate 50 L tanks in suspended baskets with constant aeration, temperature 18.4ºC ± 0.5 and a renewal rate of 50 L day-1. The animals were fed with the same diet as described above but maintaining a constant density of 168 ± 48 seeds cm^−2^. Metamorphosis of larvae took place in those tanks.

At 112 dpf one hundred individuals from each of two families, F6 (12,74 ± 0,73 mm) and F8 (12,23 ± 1,48 mm), were selected for genetic mapping and for GWAS on colour patterns, while twenty-five individuals from each of three additional families, F2 (9,82 ± 0,85 mm), F3 (12,58 ± 0,90 mm) and F7 (13,12 ± 1,67 mm), were sampled for increasing statistical power on GWAS. The shell colour pattern of each individual was recorded as outlined before, and then the shell was gently removed and stored for further analyses and photographing. The meat of each individual was fixed in pure ethanol and sent to the Genomics Platform of University of Santiago de Compostela for DNA extraction.

### 2b-RAD library construction and sequencing

DNA was extracted from the whole meat using the E.Z.N.A. E-96 mollusc DNA kit (Omega Bio-tek) following manufacturer recommendations. Library preparation followed the 2b-RAD protocol (Wang et al., 2012) with slight modifications (Maroso et al., 2018). Briefly, DNA samples were adjusted to 80 ng μL^−1^ and digested using the IIb type restriction enzyme AlfI (Thermo Fisher). As a result, the genome is cut in fragments of 36 bp of length, with the restriction enzyme recognition site in the middle. Specific adaptors, also including individual sample barcodes, were ligated and the resulting fragment amplified. After PCR purification, samples were quantified using Qubit 2.0 fluorometer (Life Technologies, Carlsbad, CA, USA) and equimolarly pooled. The pools were sequenced on a NextSeq 500 Illumina sequencer using the 50 bp single-end chemistry in the facilities of FISABIO Sequencing and Bioinformatics Service (Valencia, Spain). The two bigger families, with parents at double concentration, were multiplexed each in one run, whereas the other seventy-five samples, without parents, were multiplexed in a third run.

### Data filtering and genotyping

Raw reads were first demultiplexed according to the individual barcodes ligated during library construction. Then, reads were filtered in three consecutive steps: (i) trimmed to 36 nucleotides (length of *AlfI* generated fragments) and discarding reads below this length, (ii) removing reads without the *AlfI* recognition site in the correct position, and (iii) removing reads with uncalled nucleotides or a mean quality score below 20 in a sliding window of 9 nucleotides. Custom Perl scripts (available on demand) were used in the first two steps, and the *process_radtags* module in STACKS v2.0 (Catchen et al., 2013) was used for the latter (-c -q -w 0.25 -s 20). Bowtie 1.1.2 (Langmead et al., 2009) was used to align the filtered reads against the recently assembled reference genome of the species (Bruzos et al., unpublished), allowing three mismatches and a unique alignment (-v 3 -m 1 --sam), so reads aligning to two or more sites were discarded. The *sam* output files were converted to *bam* files and appropriately sorted to feed the *gstacks* module in STACKS, using the *marukilow* model to call variants and genotypes. The collection of putative SNPs was exported using the *populations* module.

### Linkage map construction

Files containing the putative SNPs of the two families selected for genetic mapping (F6 and F8) were processed to conform an appropriate dataset for mapping. Those SNPs not informative in the parents of each family, and those with missing data in one or both parents were filtered, as well as SNPs with extreme deviations from Mendelian segregation (P < 0.001) and those genotyped in less than 60% of the offspring.

Genotypes of retained SNPs were properly coded as Cross Pollinator (CP) cross type with unknown linkage phase to build genetic maps using JoinMap 4.1 genetic mapping software (Stam, 1993). Firstly, markers were associated to their linkage groups using the Grouping function of JoinMap based on a series of LOD scores increasing by 1, from 4.0 to 9.0. The LOD score was then selected in each family based on the number of chromosomes of the common cockle karyotype (Insua & Thiriot-Quiévreux, 1992). Markers in linkage groups with less than 10 markers and unlinked markers were excluded from further analysis. Secondly, marker ordering was performed using the Maximum Likelihood (ML) algorithm implemented in JoinMap. Default parameters were used except the chain length that was increased from 10,000 to 20,000, and multipoint estimation of recombination frequency were in general increased to ensure convergence: length of burn-in chain, 20,000; number of Monte Carlo EM cycles, 10; and chain length per EM cycles, 5,000. The *Kosambi* mapping function was used to compute centiMorgans (cM) map distances in individual female and male maps in each family. Finally, female, and male consensus maps (from both families), and a species consensus map were built using MergeMap (Wu et al., 2011).

The combination of the ML mapping algorithm implemented in JoinMap with missing data and genotyping errors, as well as the use of MergeMap to merge the parental maps and a large number of markers, can derive in inflated map lengths (see e.g. De Keyser, et al., 2010; Martínez-García et al., 2013; Peng et al., 2016). Nevertheless, ML is more powerful and robust in ordering markers in CP populations compared to Regression Mapping (RM) (Van Ooijen, 2011). To obtain the most accurate linkage maps, a mixed strategy was implemented: i) a framework genetic map was constructed using RM only with markers without missing data; ii) then, a linear regression between ML and RM distances using the same marker pairs was performed; and iii) this value was used to adjust the map distances in the ML ordered consensus maps. All maps were drawn using MapChart ver. 2.3 (Voorrips, 2002) and LinkageMapView (Ouellette et al., 2017).

### GWAS analysis

For the study of the association between phenotypic traits and SNPs across the genome, the genotypes of mapped markers in the Consensus Map were used in the two families used for mapping (F6 and F8). Further, to increase statistical power, the same mapped markers were also used in the three additional families (F2, F3 and F7), but in this case the genotypes were also filtered for minimum allele frequency (--maf 0.05) and minimum number of genotypes (--geno 0.5) using PLINK 1.9 (www.cog-genomics.org/plink/1.9; Chang et al., 2015).

A Mixed Linear Model (Yu et al., 2006) was applied to the complete dataset of filtered genotypes and the corresponding colour phenotypes catalogued as mentioned above using rMVP (Yin et al., 2021), a parallel accelerated tool for GWAS implemented in R (R Core Team, 2021). The three colour pattern traits (circle, line and stripe) were coded as binomial traits (presence / absence). The kinship matrix was previously computed following VanRaden (2008), and the EMMA method was employed to the variance components analysis (Kang et al., 2008). rMVP and the *qqman* R package (Turner, 2018) were used to plot the results. The analyses for each trait were performed in the whole dataset and within each family separately.

### Gene mining

Coding genes included in a genomic window defined around the most significant SNPs (± 500 kb) detected in GWAS for each trait were retrieved using the common cockle genome (Bruzos et al., unpublished). Gene Ontology (GO) terms were obtained for each gene with functional annotation and enriched functions at each genomic region was assessed with agriGO v2.0 (Tian et al., 2017) using a false discovery rate (FDR) of 5%.

## Results

### Colour pattern variation in common cockle European populations

European common cockle showed diverse shell colouration patterns, both within and between populations, although a predominant colour was displayed in each population (Fig. 1). Yellow (1) was the only colour detected in the population from France and was the most abundant in the populations from Denmark, Portugal (Algarve) and UK (Wales); white (2) was predominant in populations from Spain, Netherlands and Germany; brown (4) was the least frequent phenotype, only detected in one of the populations from Portugal (Algarve); and orange (6) was rather common in the other population from Portugal (Aveiro), Netherlands and UK (Plymouth) (Fig. 2). Shell patterns in the wild adults were not as marked as in the juvenile samples obtained from crosses in the hatchery (see below). Nonetheless, the presence of different colour patterns similar to those observed in the hatchery was also observed in the wild: i) a lighter coloration in the circle-shaped umbo in the population from Portugal (Algarve); ii) a darker band (black or orange) with blurred boundaries in the lateral area of the shell on the opposite side where the ligament in populations from UK (Wales, black) and Portugal (Aveiro, orange); iii) changes in the arrangement of the periostracum (protein layer) detected in the ventral margin associated with the last growth rings in all populations, excluding Denmark, where remnants of the periostracum were detected on the entire surface of the shell. Additionally, malformation of the shell that affected the last growth rings, likely related to environmental factors, were detected in individuals from France and Portugal (Algarve) (Fig. 2).

### 2b-RAD sequencing

A total of 275 samples were sequenced using 2b-RAD: the two parents and 97 offspring of F6; the two parents and 99 offspring of F8; and 25 offspring from each of F2, F3 and F7. Around ~575 million raw reads were obtained in the first two libraries for the two large families: on average ~6.9 million for parents (range from 5,635,542 to 8,343,112) and ~2.8 million for offspring (range from 3,680 to 5,991,704). After filtering, ~75% of the reads were retained and aligned to the cockle genome (Bruzos et al., unpublished). A high number of filtered reads were discarded due to mapping to two or more genomic positions (40.74%), resulting in ~200 million of high-quality aligned reads: ~3 million from each parent (range from 2,101,828 to 3,843,549) and ~900K from each offspring (range from 1,317 to 2,158,061).

The third run, with the remaining 75 samples, yielded ~275 million raw sequences (~3.6 million per offspring), and after the filtering and alignment steps, ~92 million high-quality aligned reads were retained (~1.2 million per offspring).

### Genotyping and linkage map construction

The *gstacks* module using the *marukilow* model applied to all families yielded 318,755 loci, which resulted in 85,078 polymorphic SNPs using the *populations* module. For the linkage map only F6 and F8 families were used. One individual from each family showed a low number of valid genotypes (< 30% of SNPs genotyped) and they were removed. After quality control, 7,094 and 8,439 SNPs were retained for F6 and F8, respectively. The largest number of filtered SNPs in our study was due to missing genotypes and deviations from Mendelian segregation, which can be explained by the presence of null alleles related to polymorphism in the restriction enzyme targets, as previously reported in mollusc (Saavedra & Bachère, 2006; Hollenbeck & Johnston, 2018). There were 1,329 common informative SNPs between the two families, which means that in total 14,204 SNPs were used for the construction of the genetic map.

Separate male and female maps were built for each family. To achieve an appropriate number of LGs close to the number of cockle chromosomes (n = 19), different LOD scores were explored for each genetic map (ranging between 7.0 and 9.0), resulting in 21 LGs in all maps. For the maternal map of F6, 3,514 markers were mapped for a total length of 12,572 cM, whereas the paternal map included 3,952 markers spanning 16,692 cM (Table 1). In F8, 4,698 and 4,368 markers were mapped in the maternal and paternal maps, spanning 25,017 cM and 23,266 cM, respectively. Diagrams of the four genetic maps are shown in Supplementary Figure S1, and the relationships of all mapped markers with their respective mapping positions in Supplemental Tables S1-S4. Shared markers among parental maps were used to built a single consensus map. As a result, 13,874 SNPs were assigned to 19 LGs in the final common cockle genetic map with a total length of 51,778 cM, in accordance with the 19 cockle chromosomes (Table 1 and Supplemental Tables S5-S6). The length of the maps exceeded that expected based on the genome size considering a standard relationship between physical (Mb) and genetic distance (cM) of 1.2 and a C-value of 1.37 pg (Gregory, 2022). The observed elongation of the genetic maps is the consequence of the high number of markers, several mapping families, and the limitations of the software used (JoinMap for mapping and MergeMaps for *consensing*), consistent with previous observations (Cartwright, et al., 2007; Mester, et al., 2015; Maroso, et al., 2018).

**Table 1.** Number of markers and length of each Linkage Groups (LG) in the *C. edule* female and male maps from families F6 and F8 and in the species consensus map. The numeration of LGs from the individual maps does not match the consensus map (see Suppl. Table S6 for correspondence)

| Family | F6 |  |  |  | F8 |  |  |  | Consensus |  |
| --- | --- | --- | --- | --- | --- | --- | --- | --- | --- | --- |
|  | Female |  | Male |  | Female |  | Male |  | N_Loci | Length(cM) |
| LG | N_Loci | Length(cM) | N_Loci | Length(cM) | N_Loci | Length(cM) | N_Loci | Length(cM) |  |  |
| 1 | 332 | 1136.9 | 356 | 1582.3 | 423 | 2187.4 | 402 | 2041.2 | 1151 | 3680.9 |
| 2 | 282 | 802.5 | 334 | 1349.5 | 415 | 2305.6 | 399 | 1922.4 | 1061 | 3862.2 |
| 3 | 249 | 785.8 | 329 | 1374.8 | 356 | 1921.9 | 353 | 1712.9 | 1123 | 5712.1 |
| 4 | 233 | 751.7 | 270 | 1174.9 | 331 | 1958.7 | 285 | 1342.0 | 945 | 4373.4 |
| 5 | 219 | 797.3 | 262 | 1074.2 | 306 | 1758.1 | 278 | 1335.7 | 804 | 3129.5 |
| 6 | 199 | 768.8 | 245 | 1041.7 | 277 | 1596.7 | 270 | 1445.7 | 763 | 2751.0 |
| 7 | 197 | 735.7 | 244 | 979.3 | 274 | 1374.5 | 256 | 1132.3 | 756 | 3443.3 |
| 8 | 195 | 829.3 | 239 | 834.4 | 266 | 1469.6 | 245 | 1063.3 | 749 | 2165.8 |
| 9 | 187 | 721.1 | 236 | 893.0 | 258 | 1425.9 | 214 | 1198.1 | 894 | 2761.4 |
| 10 | 179 | 707.9 | 216 | 950.3 | 235 | 1202.4 | 201 | 1504.6 | 773 | 2979.3 |
| 11 | 176 | 589.9 | 209 | 866.2 | 222 | 1180.1 | 198 | 1234.7 | 579 | 1824.3 |
| 12 | 168 | 659.4 | 202 | 696.2 | 209 | 955.1 | 198 | 964.7 | 661 | 2646.4 |
| 13 | 167 | 730.5 | 201 | 670.4 | 204 | 830.7 | 176 | 1087.7 | 643 | 2189.3 |
| 14 | 164 | 637.4 | 161 | 711.8 | 182 | 818.8 | 167 | 897.9 | 591 | 2070.5 |
| 15 | 146 | 406.1 | 148 | 687.4 | 181 | 905.6 | 161 | 976.9 | 498 | 2083.4 |
| 16 | 119 | 329.1 | 83 | 526.0 | 166 | 1014.1 | 143 | 751.6 | 627 | 2158.7 |
| 17 | 100 | 336.8 | 72 | 290.3 | 152 | 852.8 | 130 | 837.0 | 521 | 1784.7 |
| 18 | 89 | 222.9 | 47 | 329.2 | 117 | 613.4 | 110 | 611.8 | 290 | 860.0 |
| 19 | 64 | 295.7 | 43 | 321.2 | 69 | 395.9 | 87 | 661.2 | 445 | 1302.6 |
| 20 | 36 | 221.9 | 41 | 270.4 | 40 | 198.3 | 84 | 472.8 | - | - |
| 21 | 13 | 105.5 | 14 | 69.4 | 15 | 51.5 | 11 | 71.7 | - | - |
| Unlinked | 76 | - | 236 | - | 102 | - | 159 | - |  |  |
| Total | 3514 | 12572.1 | 3952 | 16692.8 | 4698 | 25017.2 | 4368 | 23266.3 | 13874 | 51778.7 |

To build a reliable framework genetic map we used the Regression Mapping approach, an approximation similar to that followed in *C. gigas* by Hedgecock et al. (2015). Accordingly, a total of 831 and 340 markers without missing data were selected in offspring of F6 and F8, respectively. Separate maps were built for each parent in each family with a LOD score ≥ 5.0. Graphical representation was not implemented but the individual groups were merged directly with the “Combine Groups” option of the JoinMap to build a consensus map. The regression of common markers distance was used to correct the distances in the original consensus map to build a new corrected-length consensus map, which included the 19 LGs but with a new total length of 1,073 cM (Figure 3). In the reduced consensus map, the estimated inter-marker distance decreased dramatically (from 4.34 to 0.13 cM), comparable to other genetic linkage maps constructed using 2b-RAD (Jiao et al., 2014; Shi et al., 2014), and the average ratio between physical and genetic distance (0.74 Mb / cM) was also much closer to the standard expected.

**Figure 3.**
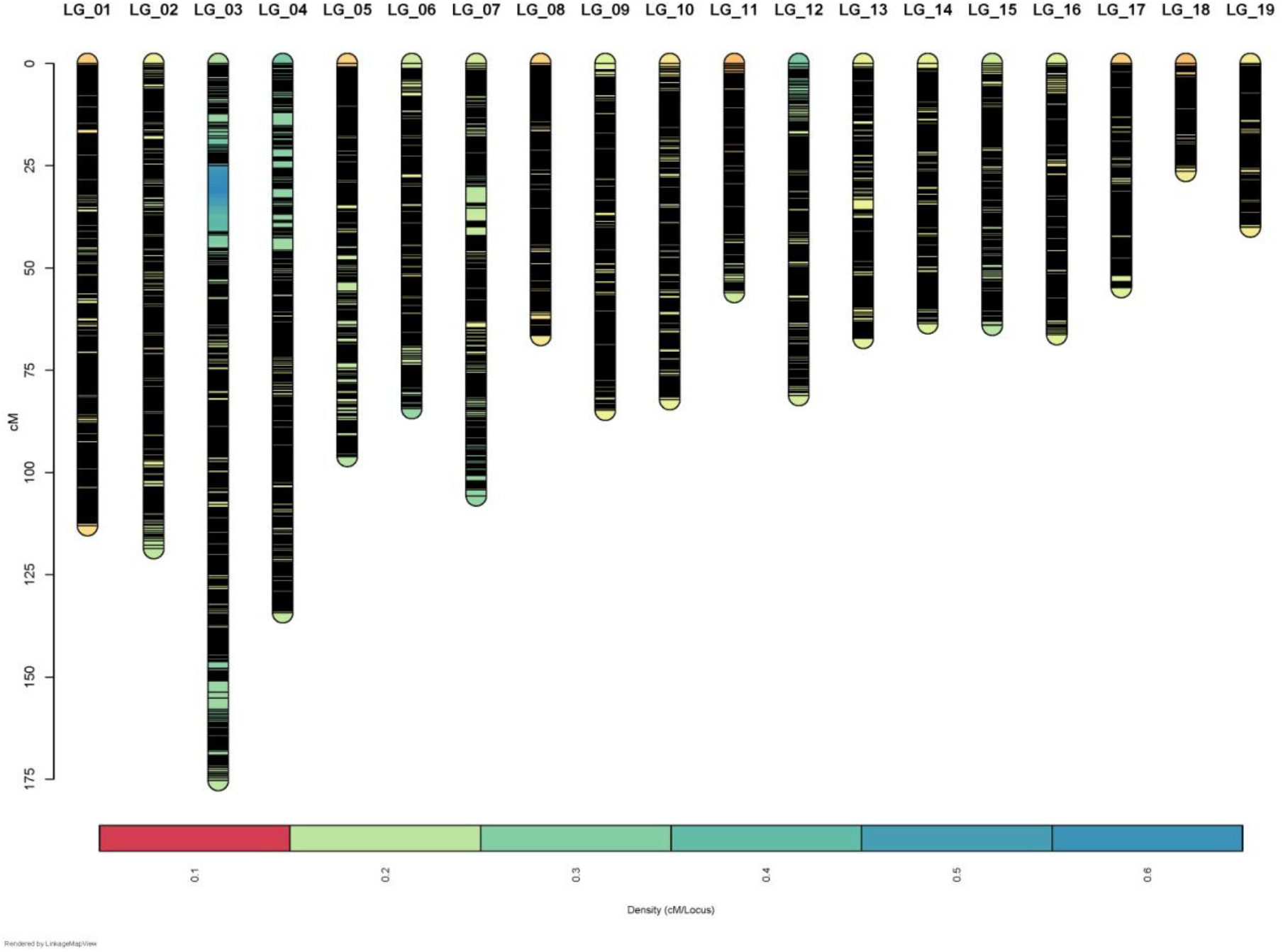
Graphical representation of the consensus map of *C. edule*. The rule in the left indicates length in centimorgans (cM).

### GWAS on shell colour patterns at hatchery

Shell colour in the five families reared in the same environmental conditions showed a remarkable variation within and between families both for the colour and the colour patterns (Fig. 4). Shell colour of the offspring of the five families was classified as outlined before (from 1 to 6) and considered as a continuous trait for analyses (Table 2). Black was the most frequent colour (31.6%), but detected only in two families (F2 and F8), whereas gray was detected only in F8, and among the other three, white was the most abundant and the only present in all families (29.8%). The orange colour was not detected in any of them, only in the wild, as outlined before. The colour patterns were only detected in three families and were quite heterogeneous; for instance the stripe was only observed in F6 at a ratio close to 1:1 (Figure 4; Table 2).

**Table 2.**
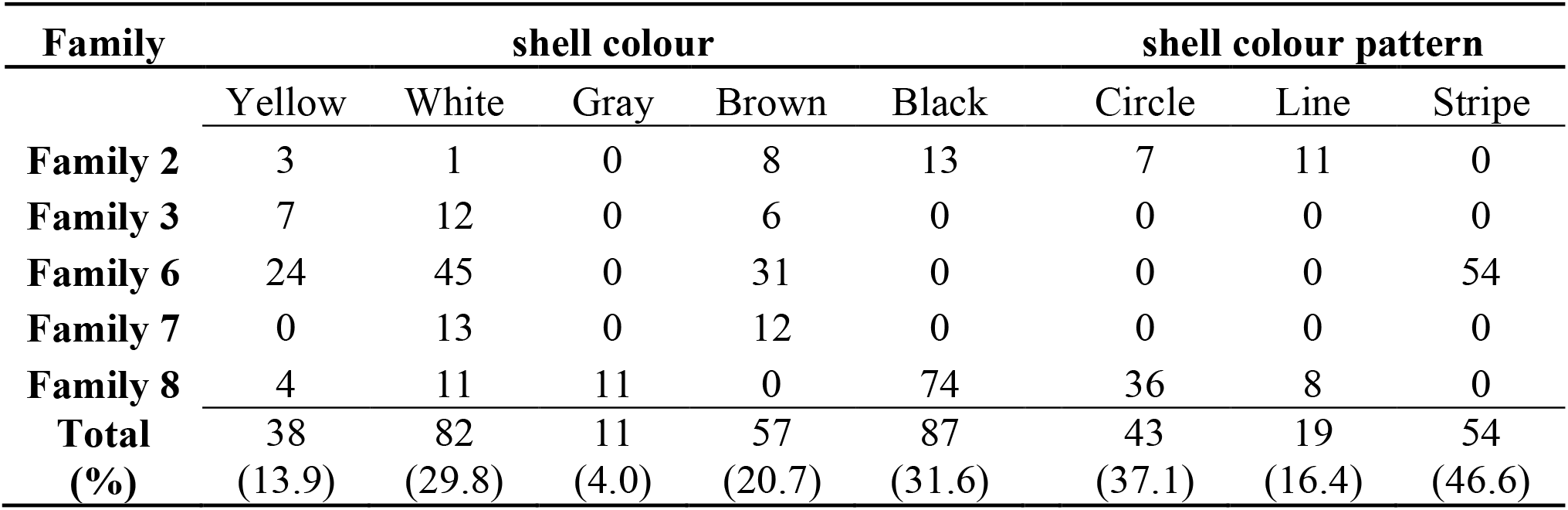
Distribution of colour phenotypes and colour patterns in the offspring of the five families of *C. edule*.

| Family | shell colour |  |  |  |  | shell colour pattern |  |  |
| --- | --- | --- | --- | --- | --- | --- | --- | --- |
|  | Yellow | White | Gray | Brown | Black | Circle | Line | Stripe |
| **Family 2** | 3 | 1 | 0 | 8 | 13 | 7 | 11 | 0 |
| **Family 3** | 7 | 12 | 0 | 6 | 0 | 0 | 0 | 0 |
| **Family 6** | 24 | 45 | 0 | 31 | 0 | 0 | 0 | 54 |
| **Family 7** | 0 | 13 | 0 | 12 | 0 | 0 | 0 | 0 |
| **Family 8** | 4 | 11 | 11 | 0 | 74 | 36 | 8 | 0 |
| **Total** | 38 | 82 | 11 | 57 | 87 | 43 | 19 | 54 |
| **(%)** | (13.9) | (29.8) | (4.0) | (20.7) | (31.6) | (37.1) | (16.4) | (46.6) |

**Figure 4.**
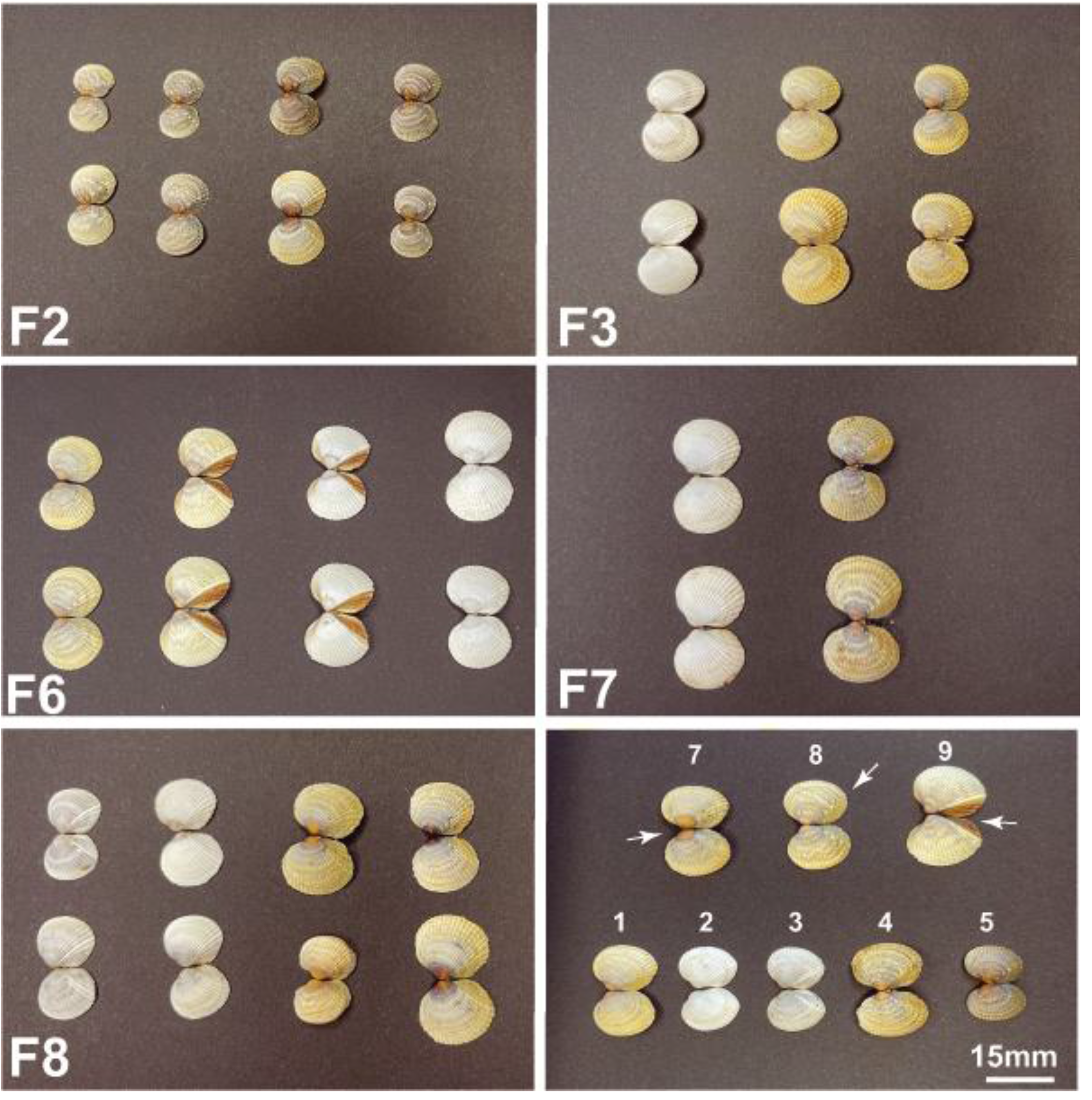
Photograph composition showing representative individuals of the five families (F2, F3, F6, F7 and F8) of *C. edule* used for GWAS on color patterns. Last panel: summary of the variation within and between families: yellow (1), white (2), gray (3), brown (4), black (5), orange (6), circle (7), line (8) and stripe (9).

The complete dataset contained colouration phenotypes for 275 individuals from five families, genotyped for 13,874 SNPs mapped in the Consensus Map (between 4,643 and 8,439 informative markers per family). GWAS revealed a highly significant genome-wide QTL for colour and stripe, and to a less extent for circle, at chromosome C13 (Figure 5). Other QTL significant at chromosome-level were detected for circle at C8, for line at C2, C9, C12 and C14, and for stripe at C1, C5, C9, and C17 (Supplemental Table S7). The estimated heritabilities were high for colour and stripe, 0.755 and 0.657, respectively, and moderate-high for circle and line, 0.537 and 0.506, respectively.

**Figure 5.**
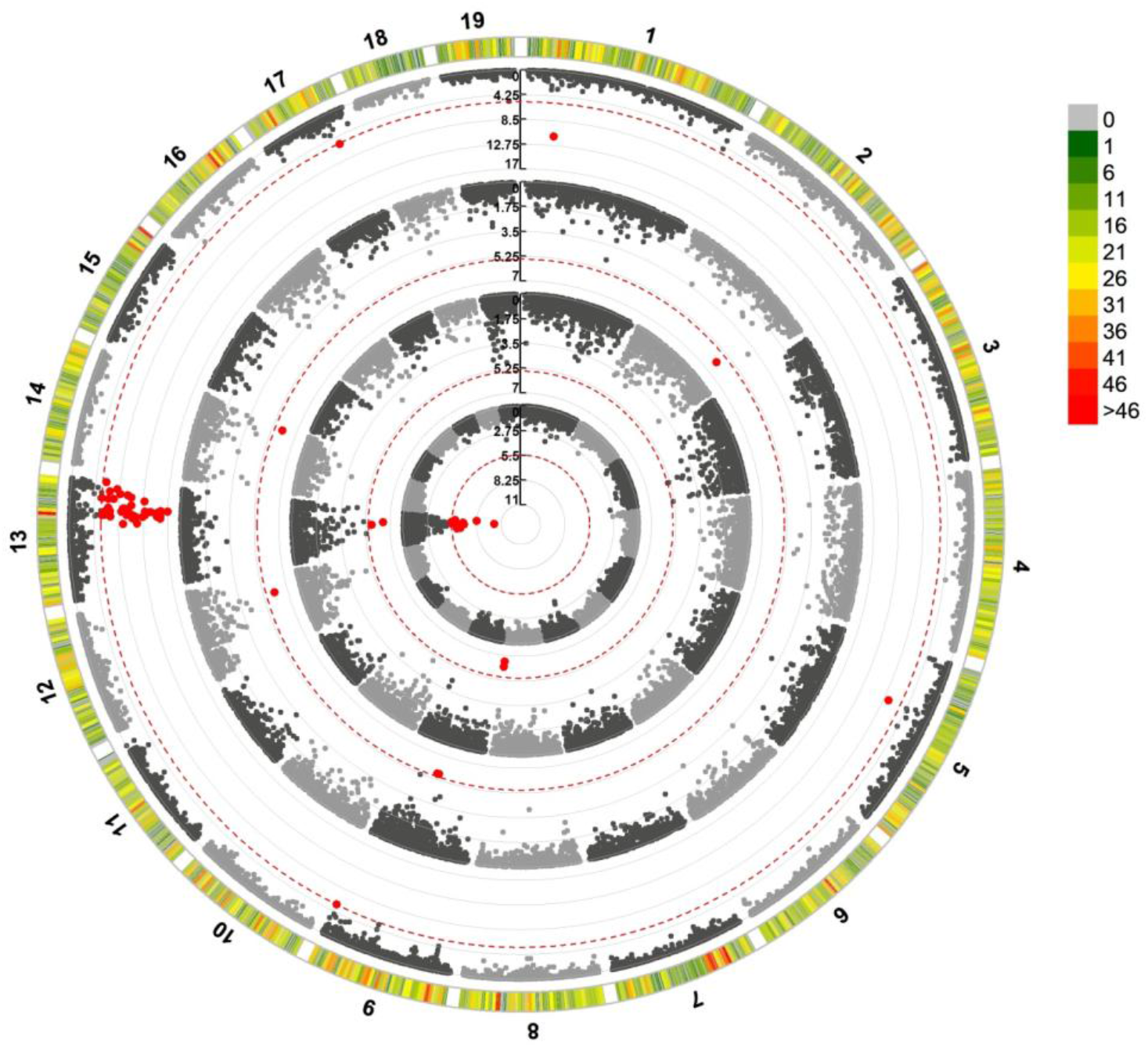
Circular Manhattan plot for SNP significantly associated with colour and colour pattern phenotypes of shell in *C. edule* based on the Mixed Linear Model (MLM). SNP-GWAS circle representations: inner-most: colour; first-middle: circle: second-middle: line; and outer-most: stripe. For Manhattan plot, Y-axis represents − log 10 (p-value) of the association with each SNP and X-axis is physical position in bp. The dashed red lines represent the Bonferroni threshold at genome-wide level (P ≤ 0.05). SNP markers are represented by gray dots and those above this threshold in red dots.

### Gene mining

According with GWAS, the most relevant region associated at genome-wide level with colour, stripe and circle was located at C13 between 14,367,847 and 33,654,270 bp (Fig. 5). The highest significant SNP associated with the three traits was located at 30,286,849 bp, but across this wide region, there were several stretches defined by highly significant SNPs associated with colour and stripe, the most significant traits (Fig. 6). Two subregions at its ends were mainly associated with colour, and mining around the most significant SNPs (14,778,145 and 33,225,111 bp ± 500 kb) disclosed 16 and 17 annotated genes, respectively (Supplementary Table S7). Half of the genes in the first window, related to shell architecture and colour, clustered on a ~250 kb region (Fig. 6; Supplemental Table S7): five were related to chitin binding (three microfibril-associated glycoprotein 4, one including a fibrinogen domain, and one DNA damage-regulated autophagy modulator), one to calcium binding (ependymin-related), one to mucin secretion, and one to ammonium transport. The second window (33,225,111 kb), included two genes related to iron binding (two steroid 17-alpha-hydroxylase/17,20 lyase-like) and another one to calcium transport (phosphatidylinositol 4,5-bisphosphate phosphodiesterase). Several Gene Ontology (GO) terms resulted enriched regarding the general cockle transcriptome (FDR < 5%), including anatomical structure development, ion transport and cell periphery (Supplemental Table S8).

**Figure 6.**
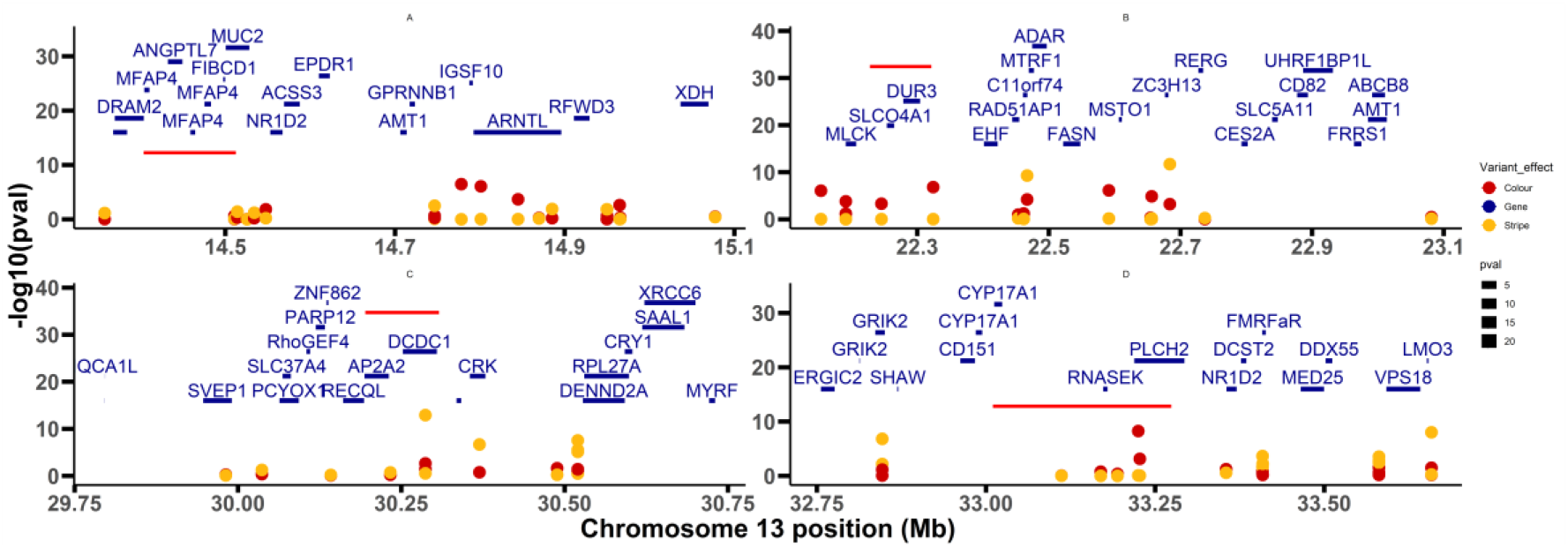
Manhattan plots for SNP significantly associated with colour (A, D) and stripe pattern (B, C) phenotypes in four chromosome 13 subregions of *C. edule* based on the Mixed Linear Model (MLM). Y-axis represents − log 10 (p-value) of the association with each SNP and Y-axis is physical position in bp. Codes of genes mined in those regions are shown with underlined bars representing their length in bp; yellow and red dots highlight associated SNPs with stripe and colour, respectively; red bars at each plot embrace the most interesting candidates in the subregion.

On the other hand, two consecutive subregions associated with stripe colour pattern were located around the most significant SNPs at 22,683,835 and 30,286,849 bp (± 500 kb) on C13 and comprised a total of 36 annotated genes. These group of genes were significantly enriched in protein binding and active transmembrane transporter activity GO terms and included three cell membrane organic transporters (solute carrier organic anion transporter family member 4A1-like, urea-proton symporter DUR3-like and sodium/glucose cotransporter 4-like); one chitin-binding (sushi, von Willebrand factor type A, EGF and pentraxin domain-containing protein 1-like); several involved in transit across the membrane by endo-exocytosis mechanisms (prenylcysteine oxidase 1-like, glucose-6-phosphate exchanger SLC37A4-like, and AP-2 complex subunit alpha-2-like); and one related to iron chelation (putative ferric-chelate reductase 1).

Other chromosome-level significant SNPs were located at C1 (9,307,825 bp), C5 (17,329,873 bp), C9 (38,779,506bp) and C17 (21,420,081bp), related to stripe; at C8 related with circle (38,336,724bp); and at C2 (44,887,010bp), C9 (22.021.180bp), C12 (27,494,091bp) and C14 (31,638,389bp), related to line (Supplemental Table S7).

## Discussion

Molluscs represent a highly diverse Phylum of invertebrates comprising an estimated number of 200,000 species, distributed across almost every type of habitat worldwide (Gomes dos Santos et al., 2020). They have key roles as ecosystem engineers, water filtering and pollution monitoring, jewelry, and, of course, as an important food source. More than 17 million metrics tons of molluscs were farmed worldwide in 2018 and most of this production concentrated in a handful of species of the class Bivalvia (FAO, 2022). Bivalve aquaculture mainly relies on extensive farming based on the collection of wild seed and harvesting in natural beds, which means that wild populations are under important human alterations (Hollenbeck & Johnston, 2018).

Shell constitutes a main structure of mollusc anatomy that protects them against predators and desiccation, but that also plays other important functions depending on species. Shells are secreted by the mantle, so colour patterns are mainly determined by pigments produced by this tissue, although the microstructure of the shell may also contribute to coloration (Clark et al., 2020; Saenko & Schilthuizen, 2021). Despite the efforts made in recent years, there is still a knowledge gap regarding the genetics underpinning shell colour in Mollusca and its potential role for adaption. While it is well-known that shell colour can be under genetic control, biotic and abiotic factors, such as diet, temperature, salinity or pH, also play a role, and the interaction between them is not well understood yet (Williams, 2017). To address the genetic architecture of complex traits, such as shell colour pattern, comprehensive genomics approaches are essential, and although mollusc genomic resources have lagged behind those of vertebrates, today their application is affordable even in species with scarce genomic resources (Liu et al., 2021).

Genomic resources have increased exponentially in the last decade as a consequence of the lowering cost of sequencing technologies and the new bioinformatic tools that enabled the assembling of genomes at chromosome level, the construction of high-density genetic maps and the genotyping millions of SNPs for genomic screening (Xu et al., 2020). Genetic maps are essential for the identification of genomic regions underlying phenotypic variation for relevant traits under domestic or natural selection (Saavedra & Bachère, 2006; Hollenbeck & Johnston, 2018; Tan et al., 2020). The first mollusc genetic maps were published in *Crassostrea virginica* (Yu & Guo, 2003) and *Crassostrea gigas* (Hubert & Hedgecock, 2004; Li & Guo, 2004) using AFLPs and microsatellites, respectively, but the lowering cost of sequencing technologies enabled genotyping thousands of SNPs for improving map density (Nie et al., 2017; Li et al., 2018). The common cockle genetic map here constructed comprehends 13,874 markers with an inter-marker distance of 0.16 cM and comprises the 19 expected LGs matching with the haploid karyotype of the species (Insua & Thiriot-Quiévreux, 1991), being, to our knowledge, the denser genetic map published to date in molluscs. Besides its importance for genomic screening, the common cockle genetic map is an invaluable tool for genome scaffolding, as has been previously reported in other species (Martínez et al., 2021; de la Herrán et al., 2022), and in fact, five of the seven mollusc genomes assembled at chromosome-level took advantage of high-density linkage maps (Hollenbeck & Johnston, 2018).

We used the common cockle genetic map to ascertain the genetic component underlying the broad diversity observed for colour pattern on the wild populations of this species in the Northeast Atlantic. Ricardo et al. (2017, 2022) showed an important variation in shell ion composition in common cockle, apparently related to environmental factors, that made possible to trace back the geographic origin of specimens, but they did not associate this variation with colour. Most phenotypes observed in wild populations in our study could be identified in the families produced at hatchery, although their intensity was somewhat faded, likely due to environmental factors operating across the lifespan of the adult specimens collected. Indeed, differentiation of individuals by colour patterns in families was very clear, sometimes resembling single gene Mendelian segregation, and furthermore, heritabilities for all colour traits evaluated were high (h^2^ > 0.5), supporting a significant genetic component underlying colour variation in common cockle. The fact that most phenotypes observed in the Atlantic distribution appear to be segregating in a single population from NW Spain supports an important intrapopulation variation in common cockle and suggest that more detailed studies across the full lifespan could disclose more variation than observed in our preliminary screening.

In accordance with these observations, the GWAS performed on five full-sib families produced at hatchery identified a major QTL at chromosome 13 explaining most of the variance observed for two of the four traits evaluated (colour and stripe). This region encompassed ~13 Mb, although with different stretches for the same or the different traits studied, which suggests the existence of a broad gene cluster related to colour pigmentation in common cockle with different genes playing diverse functions on similar traits. A total of 69 annotated genes involving several enriched GO terms, including anatomical structure development, ion transport, transmembrane transporter activity, protein binding and cell periphery, were found on this region after mining the cockle genome (Bruzos et al., unpublished). We identified a notable proportion of genes related to ion binding and transport/secretion across the cell membrane, such as ammonium or organic transporters, calcium binding or iron chelation, mucin production and several related to endo-exocytosis mechanisms, which play an important role in the development of the shell and that has been related with shell colour in other molluscs (Ding et al. 2015; Teng et al., 2018; Wang et al. 2018; Hu et al., 2019; Liao et al., 2019; Xu et al. 2019). Moreover, shells are secreted by the mantle in a process called biomineralization, where chitin represents an important component (Schönitzer & Weiss, 2007; Furuhashi et al., 2009; Lemer et al. 2015). We could identify six chitin-binding related genes, four of them in a small genomic region encompassing ~110 kb including three microfibril-associated glycoprotein 4 genes. Genes related to chitin and calcium metabolism involved in different shell colour lines have been reported in *Pecten yessoensis* (Ding et al., 2015). Nevertheless, it is important to note that previous studies have shown that some of the genes related to shell architecture and colour are species-specific (Williams, 2017; Saenko & Schilthuizen, 2021), so further studies, should be conducted in the future to ascertain the roles of the genes located at the C13 cluster for a deep understanding of colour pattern diversity in common cockle.

A final reflection on the putative adaptive role of colour pattern diversity in the common cockle is worth a final thought. While shell colour can be important for the adaptation of bivalve populations to selective pressures, this might not be a major factor in molluscs that live buried in the sediment, such as *C. edule*. Nevertheless, cockles live in the intertidal zone and constitute the feed for different predators, such as birds, mammals and crustaceans, among others. Moreover, a broad colour pattern diversity exist in this species across its full distribution range and, importantly, with a substantial underlying genetic variation, as the high heritabilities estimated in our study demonstrate, so its putative adaptive role should deserved further studies. Interestingly, the C13 colour associated QTL overlaps with a genomic region in the same chromosome which showed very consistent signals of divergent selection, including several outliers above the neutral background and highly significant linkage disequilibrium suggestive of selective sweeping (Vera et al., 2022).

## Conclusion

Here we presented a high-density genetic map in common cockle, the first reported in the species and, to our knowledge, the highest dense map reported in molluscs to date. The consistency of the map was shown by fitting the number of linkage groups to the haploid chromosome number of the species and by the consistent result of the GWAS on colour pattern. This map was used to ascertain the genetic component and architecture underlying the broad colour pattern diversity observed in common cockle in the Northeast Atlantic. High heritabilities in all traits evaluated support an important genetic component, which apparently was mostly related to a single genomic region at C13, where a cluster of genes related to specific enriched functions related to shell architecture and colour were detected. Our results suggest a potential adaptive role of the variation observed and highlights the importance of deeper studies at population level across the cockle lifespan to understand the significance of the variation observed.

## Acknowledgements

The research leading to these results has received funding from the Interreg Atlantic Area Programme through the European Regional Development Fund for the project Co-Operation for Restoring CocKle SheLl fisheries and its Ecosystem Services in the Atlantic Area (COCKLES, EAPA_458/2016; www.cockles-project.eu). Authors wish to thank L. Insua, S. Sánchez-Darriba for their technical support. Alicia L Bruzos was supported by a predoctoral fellowship from the Spanish Ministry of Economy, Industry and Competitiveness (BES2016/078166). SCUBA CANCERS is funded by European Research Council (ERC) Starting Grant 716290 of Jose Tubio. Bioinformatic analysis were supported by Centro de Supercomputación de Galicia (CESGA).

## Author Contributions

M.H conducted all analysis and drafted the manuscript. D.R. collaborated in the analysis and the experiment design. D.C. performed the experimental crosses. S.D were involved in the experimental crosses, sampling, laboratory analysis and phenotyping. A.L.B was involved in sampling and gene mining. A.B designed custom scripts and supervised the bioinformatic analysis. P.M. conceived the study, supervised the project and revised the manuscript. All authors collaborated in the manuscript and approved the final version.

## Legends to Supplementary Figure

**Supplemental Figure S1**. Graphical representation of maternal and paternal genetic map of the two mapping families of *C*.*edule*. The rule in the left indicates length in centimorgans (cM).

## Notes

### Competing Interest Statement

The authors have declared no competing interest.

