## Supplementary figures and images for "The broad shell colour variation in common cockle (*Cerastoderma edule*) from Northeast Atlantic relies on a major QTL revealed by GWAS using a new high-density genetic map"

### Supplemental Fig. S1

#### Slide 1
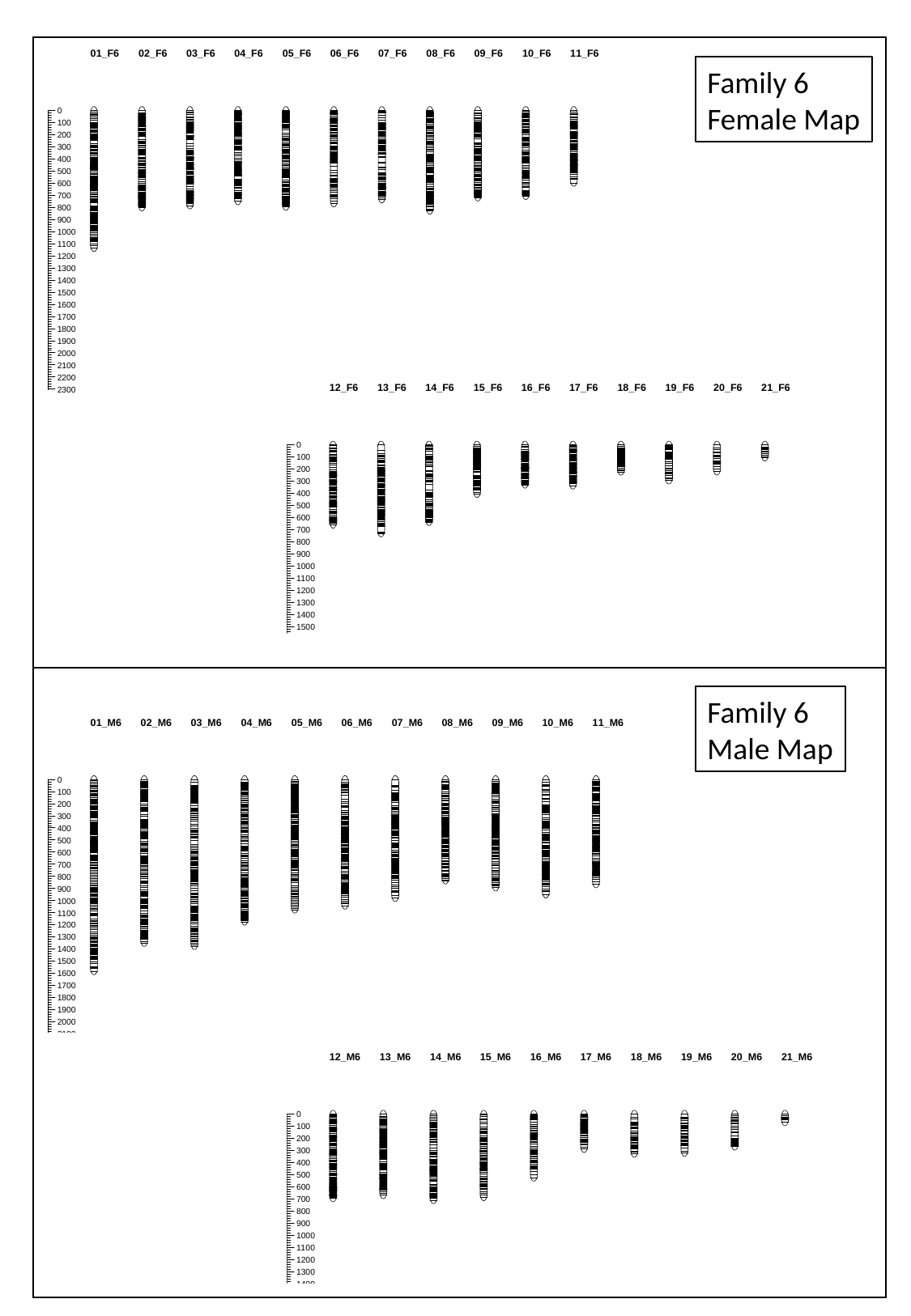

Family 6
Female Map
Family 6
Male Map

#### Slide 2
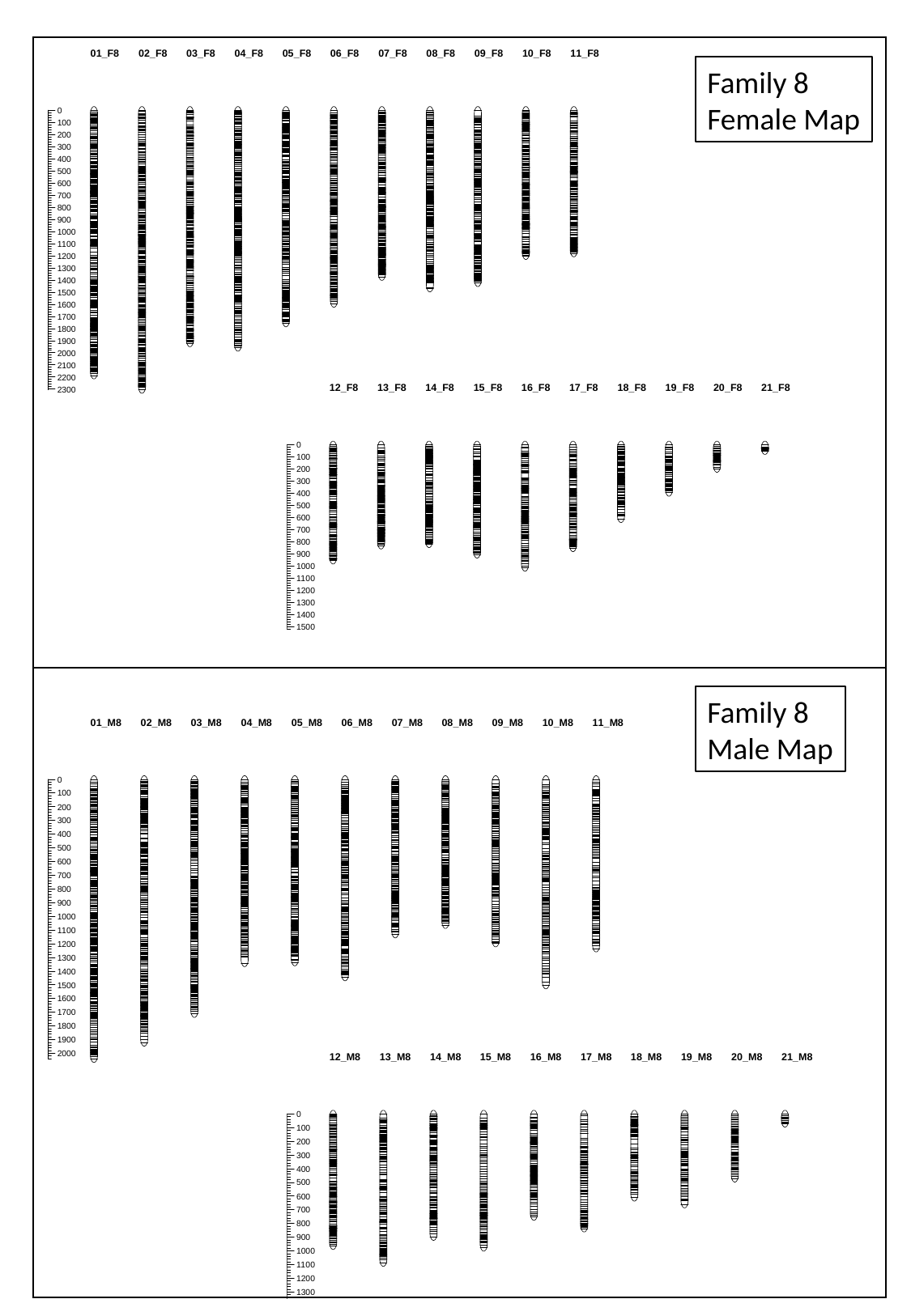

Family 8
Female Map
Family 8
Male Map
